# Towards A More Informative Representation of the Fetal-Neonatal Brain Connectome using Variational Autoencoder

**DOI:** 10.1101/2022.06.16.496454

**Authors:** Jung-Hoon Kim, Josepheen De Asis-Cruz, Dhineshvikram Krishnamurthy, Catherine Limperopoulos

## Abstract

Recent advances in functional magnetic resonance imaging (fMRI) have helped elucidate previously inaccessible trajectories of early-life prenatal and neonatal brain development. To date, the interpretation of fetal-neonatal fMRI data has relied on linear analytic models, akin to adult neuroimaging data. However, unlike the adult brain, the fetal and newborn brain develops extraordinarily rapidly, far outpacing any other brain development period across the lifespan. Consequently, conventional linear computational models may not adequately capture these accelerated and complex neurodevelopmental trajectories during this critical period of brain development along the prenatal-neonatal continuum. To obtain a nuanced understanding of fetal-neonatal brain development, including non-linear growth, for the first time, we developed quantitative, systems-wide representations of neuronal circuitry in a large sample (>700) of fetuses, preterm, and full-term neonates using an unsupervised deep generative model called Variational Autoencoder (VAE), a model previously shown to be superior to linear models in representing complex resting state data in healthy adults. Here, we demonstrated that non-linear brain features, i.e., latent variables, derived with the VAE, carried important individual neural signatures, leading to improved representation of prenatal-neonatal brain maturational patterns and more accurate and stable age prediction compared to linear models. Using the VAE decoder, we also revealed distinct functional brain networks spanning the sensory and default mode networks. Using the VAE, we are able to reliably capture and quantify complex, non-linear fetal-neonatal functional neural connectivity. This will lay the critical foundation for detailed mapping of healthy and aberrant functional brain signatures that have their origins in fetal life.

## 1. Introduction

In-utero fetal brain development follows a highly organized, dynamic and precisely patterned process that lays a critical foundation for lifelong neurodevelopmental and neuropsychiatric brain health^1, 2^. The architecture of the human connectome also changes rapidly with brain maturation and plays a critical role in critical periods of early brain development^3, 4^.

Functional magnetic resonance imaging (fMRI) has become a powerful neuroimaging tool for investigating whole brain dynamics^5^. Seminal fMRI studies have shown that the mature human brain is comprised of several overlapping large-scale brain networks subserving different sensorimotor and higher cognitive functions^6, 7^. These functional brain networks (FBNs) evolve over the life span, from infancy to old age^8, 9, 10, 11^. The recent successful application of resting-state fMRI to the human fetus and newborn has extended these critical observations and shed new light on the emergence of these FBNs in the first 1000 days of life. Sensorimotor networks have been shown to develop around the third trimester of pregnancy ∼30 weeks^12^, while higher cognitive networks emerge later in neurodevelopment^13^. For example, the primitive form of the default mode network emerges around 35 weeks in fetal^12, 14^ or newborn^15^ resting-state fMRI (rsfMRI) and highly resembles the adult configuration by the end of the first year^11^. However, to date, analysis of fetal and neonatal rsfMRI data has heavily relied on linear models, using heuristic knowledge learned from adult rsfMRI studies such as brain atlases^13^ brain network patterns^16^, or driven by data-driven linear computational models such as independent component analysis (ICA)^17^. While modeling adult rsfMRI activity as a linear system has been shown to be a simple and reasonable approach^18, 19, 20^ (see review^21^), the utility of linear models in fetal-neonatal fMRI analysis can be limited due to the rapid neurodevelopment occurring during the fetal-neonatal stage^22, 23, 24^.

It is now recognized that the hemodynamic function linking neural activity to the measured fMRI signal is non-linear^25, 26, 27^. Consequently, recent studies with non-linear modeling of fMRI data have yielded more informative brain representations compared to conventional linear methods, leading to significant improvements in disease classification^28^ and identifying individual-specific (i.e., as opposed to group-averaged) brain features based on functional connectivity (FC)^29^, among others. In addition to the non-linearity inherent in fMRI measurement, studies have shown that regions of the brain mature at different rates^15, 17, 30^. For example, fetal inter-hemispheric long-range connectivity has an inflection point between 26^-^29 weeks, when connectivity strength rapidly increased in a region-specific manner (i.e., occipital before temporal before frontal region), and then plateaus at around 32 weeks^31^. Taken together, these findings suggest that longitudinal investigation of fetal-neonatal FBNs using rsfMRI likely necessitates a more comprehensive computational approach that encompasses and models possible non-linearity of fetal and neonatal rsfMRI data. To the best of our knowledge, most investigations of emerging FC have utilized linear models, which may fail to capture important non-linear information of the highly dynamic fetal-neonatal brain continuum that is critical for more accurate characterization of functional brain development in the first nine months of life.

Recently, we developed a novel analysis framework for adult rsfMRI using unsupervised deep generative model – Variational Autoencoder (VAE)^29^. Our VAE model was able to learn a non-linear feature set (or “latent space”) effectively using large-scale adult rsfMRI data. Our preliminary results in adults^29^ demonstrated that the fully trained VAE model could disentangle generative factors of rsfMRI data and encode the learned representations as latent variables. Noteworthy, generated VAE representations in the latent space were robust over varying signal quality of rsfMRI. This initial success motivated us to apply the VAE model to characterize in-vivo fetal-neonatal brain maturation, and to critically investigate the non-linearity of fetal-neonatal rsfMRI data. We hypothesized that non-linear representations of fetal-neonatal rsfMRI, learned through the VAE model, would carry more accurate and informative neural signatures of the prenatal-neonatal brain maturational continuum, compared to linear representations defined at the network scale using ICA or at the regional scale using multi-modal cortical parcel. To validate our hypothesis, we conducted two experiments: first, compression of instantaneous fMRI patterns of subjects and second, prediction of gestational/postmenstrual ages using their rsfMRI scans. The generalizability of our findings was carefully tested using two large datasets acquired by different institutions, together consisting of >500 fetuses, premature and healthy term-born neonates. Finally, using latent variables at the group-level, we mapped fetal-neonatal brain resting state networks at different age groups.

## 3. Results

### 2.1. VAE represented fetal-neonatal fMRI patterns better than linear models

We first determined whether our VAE model pretrained using adult rsfMRI data could extract meaningful representations of fetal-neonatal cortical activity. To accomplish this, we measured the reconstruction performance of rsfMRI patterns via the VAE model and compared it to linear representations derived using ICA and cortical parcels. For the latter, we employed linear latent spaces defined through group ICA of adult rsfMRI, the same data set used for VAE training. As the number of dimensions of latent space (i.e., number of features) is critical to reconstruction performance, we utilized linear latent spaces with different dimensions (i.e., IC50, IC100, IC200, and IC300; publicly available at https://db.humanconnectome.org/ ^32^). Reconstruction performance was defined as the spatial correlation between original and reconstructed cortical patterns (Fig. 1).

**Figure 1.**
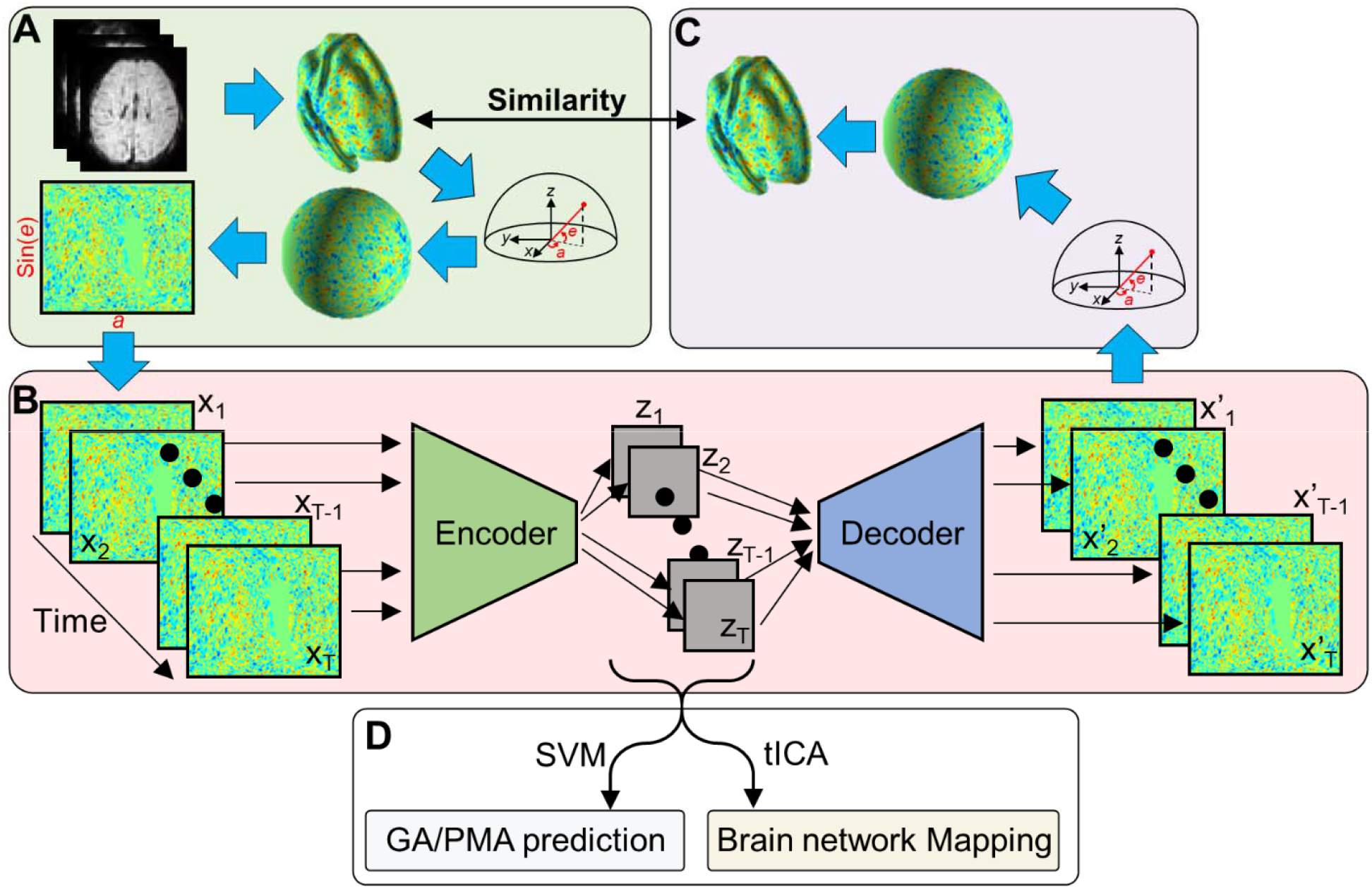
Variational Autoencoder (VAE) and its application on fetal-neonatal fMRI data. **(A)** From volumetric fMRI patterns, 2D brain pattern is estimated via geometric reformatting. **(B)** Through the encoder and decoder of VAE, each brain pattern is compressed to latent variable z and reconstructed to x’. **(C)** Reconstructed 2D image is re-shaped into cortical space through the inverse reformatting step. **(D)** Latent representations estimated by VAE are used as features of different analysis.

Qualitatively, we observed that the VAE model successfully reconstructed the fetal-neonatal cortical patterns but at the cost of some smoothing effects (Fig. 2. A and B). While linear latent spaces with different dimensions 50, 100, 200, and 300 (IC50-300) also exhibited reasonable reconstruction performance, VAE showed more accurate reconstruction patterns compared to the original image, especially in the frontal (Fig. 2. A) and temporal regions (Fig. 2. B). To confirm our qualitative observation, we quantified the spatial similarity between original and reconstructed brain activity patterns, for each time point (Fig. 2. C and D). Although spatial similarity varied over time course, VAE always showed the best similarity compared to other linear counterparts. This observation remained consistent at the group level and across different datasets (Fig. 2. E). Interestingly, IC maps with higher dimensions (=300) than VAE (=256) showed inferior reconstruction performance than VAE model (Fig. 2. E). We further confirmed that the difference in reconstruction performance between VAE and IC300 was statistically significant (paired t-test, Bonferroni-corrected *p*<10^−3^), for all datasets. Collectively, these results suggest that the non-linear latent representation obtained with VAE delineated the spatiotemporal characteristics of fetal-neonatal rsfMRI more precisely than its linear counterparts.

**Figure 2.**
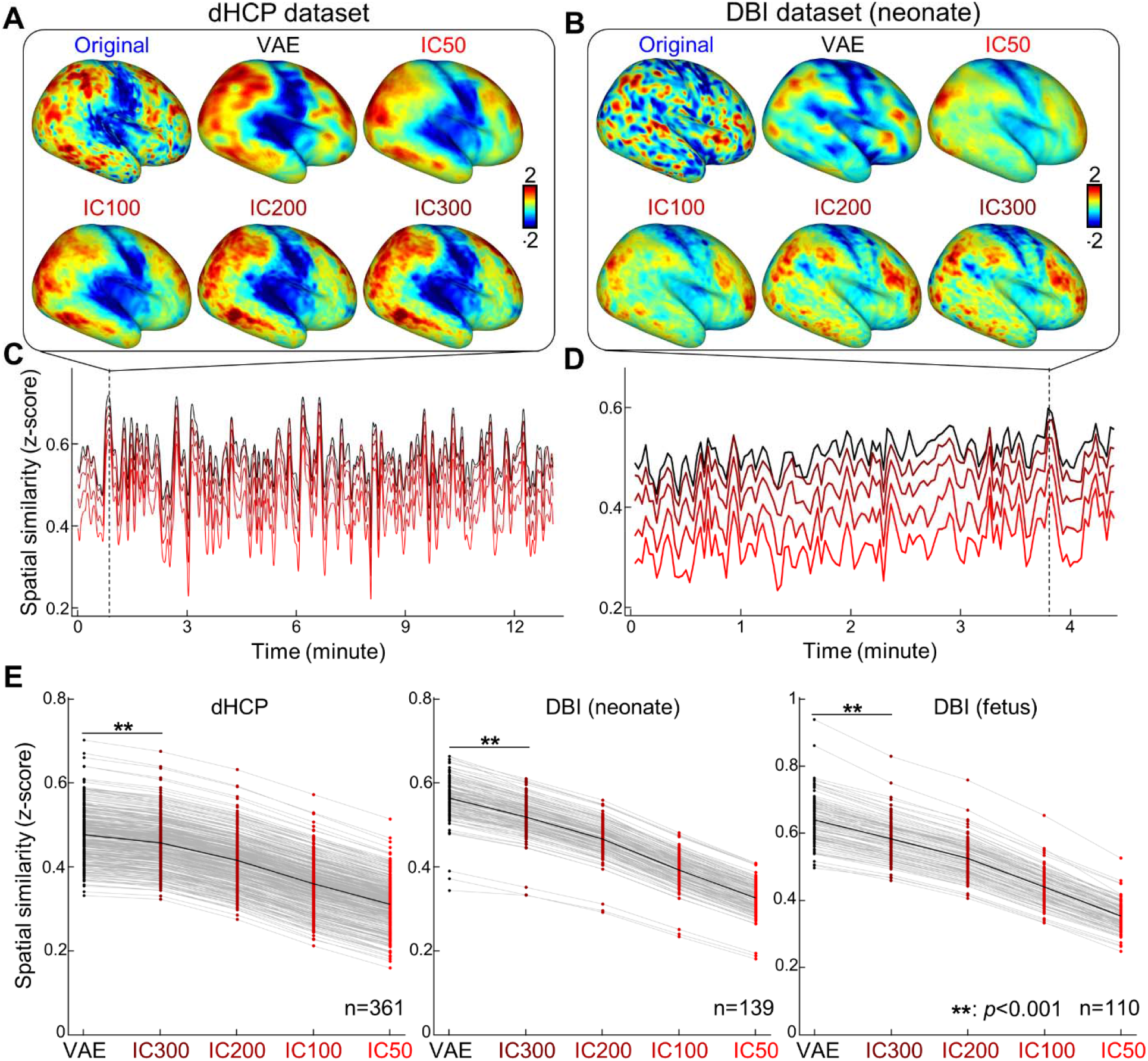
VAE represented fetal-neonatal fMRI patterns better than linear counterparts. Compared to linear spaces defined by group independent component analysis (IC50, IC100, IC200, and IC300), VAE shows the best reconstruction performance on both dHCP **(A, C)** and DBI datasets **(B, D)**, at the individual level and at the group level **(E). \*\***: *p*<0.001, Bonferroni-corrected.

### 2.2. Reconstructed fMRI patterns contained not only age, but also individual neural signatures

We speculated that a critical factor of subject-wise variation in the reconstruction performance was the variability in participants’ ages at the time of the MRI study. We found that post-menstrual age (PMA) at MRI was significantly and positively correlated with the reconstruction performance, for both dHCP (Fig. 3. A; *r*=0.20, *p*<0.01) and DBI (Fig. 3. B; *r*=0.35, *p*<0.01) datasets. Conversely, for the fetal dataset, we found that the reconstruction performance was negatively correlated with GA (Fig. S1. A; *r*=-0.59, *p*<0.01). We posit that this was in part due to the smoothing effect induced when registering the small size of the fetal brain to the standard 40-week neonatal brain template (see Supplemental Information for details).

**Figure 3.**
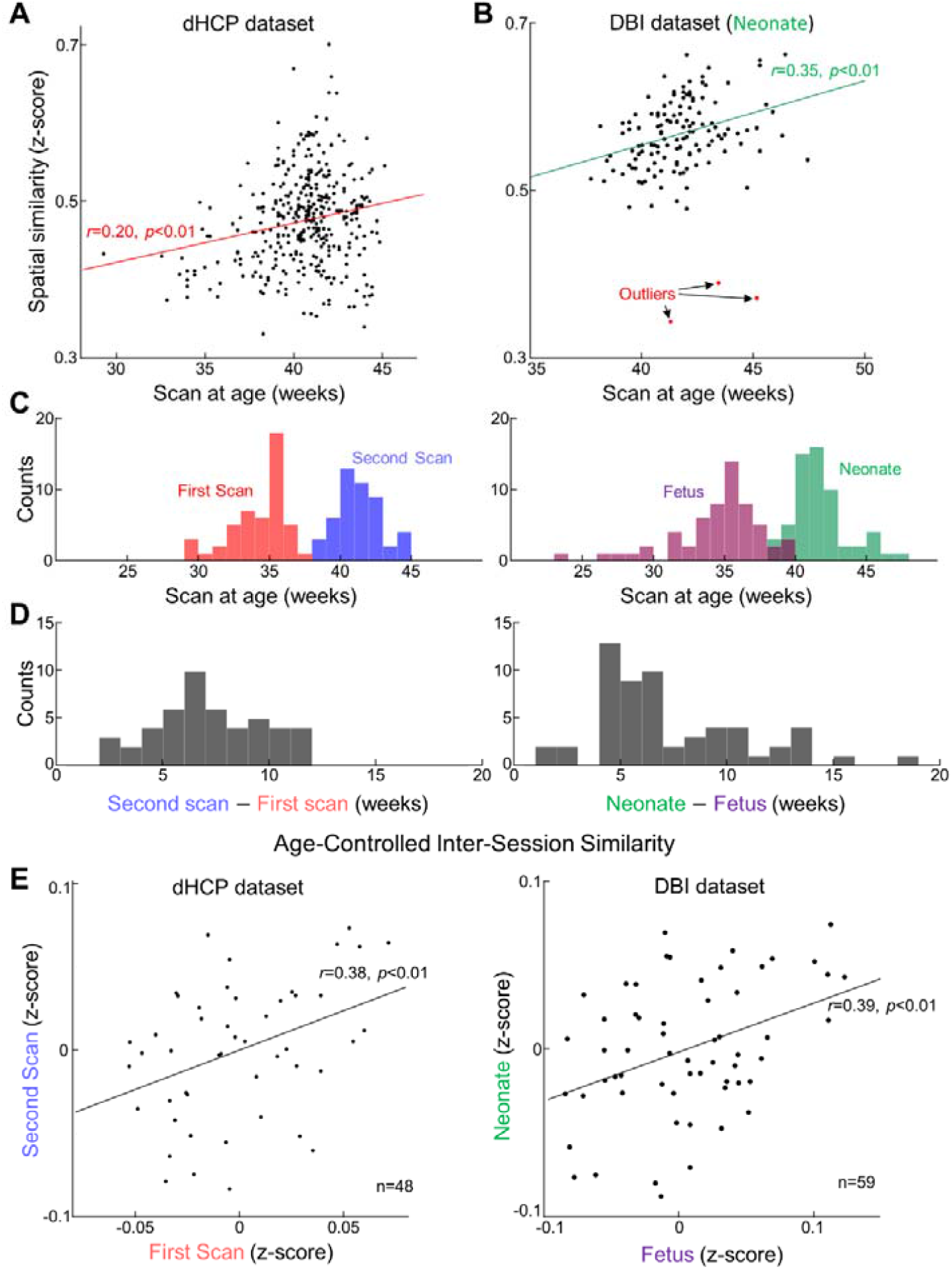
Reconstruction degree of fMRI patterns not only varies across different ages at scan but also is individual-specific. Reconstruction degrees across individuals are positively correlated with their ages at scan for both datasets, dHCP **(A** and DBI **(B)**. In **(B)**, each red dot stands for the outliers having > 3 median absolute deviations away from the median. **(C)** Distributions of ages at repeated scans (left; dHCP dataset, first and second scans; right; DBI dataset, fetal and neonatal scans). **(D)** Distributions of inter-scan age difference, for dHCP (left) and DBI dataset (right). **(E)** Scatter plot of reconstruction degree between the first scan and (or fetal scan for DBI) and the second scan (or neonatal scan for DBI), after controlling the effect from ages at scan.

Interestingly, we found that the age-dependency of the reconstruction performance was significant, but the strength of correlation was moderate *r*=0.2 (*p*<0.01) and *r*=0.35 (*p*<0.01), in the dHCP and DBI datasets, respectively. We sought to determine whether reconstruction performance in newborns was consistent across scans. To investigate this, we analyzed the reconstruction performance in subjects with two postnatal scans (dHCP dataset; n= 48 subjects; Fig. 3. C left) or those with *in-* and *ex-utero* scans (DBI dataset; n= 59 subjects; Fig. 3. C right). The time interval between scans for the dHCP dataset was 3-12 weeks (Fig. 3. D left) and 2-19 weeks (Fig. 3. D right) for the DBI datasets. After regressing out the age effect from the reconstruction performance, we found that subjects having better reconstruction performance on their first scan also tended to have better reconstruction performance on their second scan (Fig. 3. E left). This trend was also observed in subjects with in-utero and ex-utero scans (Fig. 3. E right). To confirm that this finding was not driven by similar head motion profiles between scans, we analyzed whether head motion was related between the subjects’ two scans. We found that there was no significant correlation in the inter-session frame-wise head displacement (Fig. S2 left; *r*=0.15, *p*=0.32). The inter-session similarity of reconstruction performance also remained significant after controlling for head motion (Fig. S2 middle; *r*=0.30, *p*<0.05). We further found the inter-session similarity of reconstruction performance was not driven by the variation in brain sizes (Fig. S1. F and Fig. S2 right) Altogether, our results suggest that the individual variation of the reconstruction performance of resting-state fetal-neonatal brain activity was tightly connected not only to chronological brain maturity but also to the intrinsic neural traits of the subjects even during the *in-utero* period.

### 2.3. Prediction of gestational age and postmenstrual age using VAE-derived representations

Next, we investigated whether latent variables extracted by VAE contained neural signatures of fetal-neonatal functional brain maturation using the dHCP dataset. Given different types of latent variables that were defined by VAE, cortical parcels, or IC50-300, we calculated functional connectivity patterns, which was defined by the covariance (or correlation for cortical parcels and GICA maps) for every pair of latent variables, per subject (Fig. 4. A). Through fusing 10-fold cross validation scheme and linear regression support vector machine in dHCP dataset (RSVM), we found that latent variables derived by VAE yielded highly reliable prediction accuracy (Fig. 4. B and C). As suggested in brain age prediction studies^33^, we also adjusted for prediction bias in the model (Fig. 4. C). Interestingly, the age prediction model with VAE-derived representations reached the best age predictability; mean ± standard deviation over 10 folds, root-mean-squared-error, RMSE=1.92±0.39, mean absolute error, MAE=1.48±0.27, *r*^2^=0.73±0.10, with a big margin compared to models based on linear latent features (Table 1). It is noteworthy that different from the monotonic improvement of the reconstruction performance over increasing IC #, the age predictability reached peak at IC200 while predictability deteriorated in the prediction model with features from 300 ICs (MAE: 1.81±0.22 for IC200 vs. 1.93±0.26). The age predictability of the cortical parcels (MAE: 1.87±0.30) model was between the IC300 and IC200 models. Collectively, these results suggest that non-linear features extracted by VAE improved age prediction, outperforming linear features derived using ICA or cortical parcels.

**Table 1.** Comparison of age prediction performance in dHCP dataset using different latent representations. RMSE: root mean squared error, MAE: mean absolute error, R^2^: explained variance. Red highlight indicates the best performance among different latent representation methods.

| Representation Method | RMSE (mean±SD) | MAE | R<sup>2</sup> | Correlation |
| --- | --- | --- | --- | --- |
| VAE (N=256) | 1.92 ± 0.39 | 1.48 ± 0.27 | 0.73 ± 0.10 | 0.85 |
| Cortical Parcel (N=360) | 2.36 ± 0.36 | 1.87 ± 0.30 | 0.65 ± 0.09 | 0.80 |
| IC50 | 2.86 ± 0.81 | 2.20 ± 0.56 | 0.58 ± 0.10 | 0.73 |
| IC100 | 2.61 ± 0.43 | 2.10 ± 0.38 | 0.59 ± 0.13 | 0.76 |
| IC200 | 2.31 ± 0.34 | 1.81 ± 0.22 | 0.66 ± 0.11 | 0.81 |
| IC300 | 2.42 ± 0.32 | 1.93 ± 0.26 | 0.62 ± 0.10 | 0.78 |

**Figure 4.**
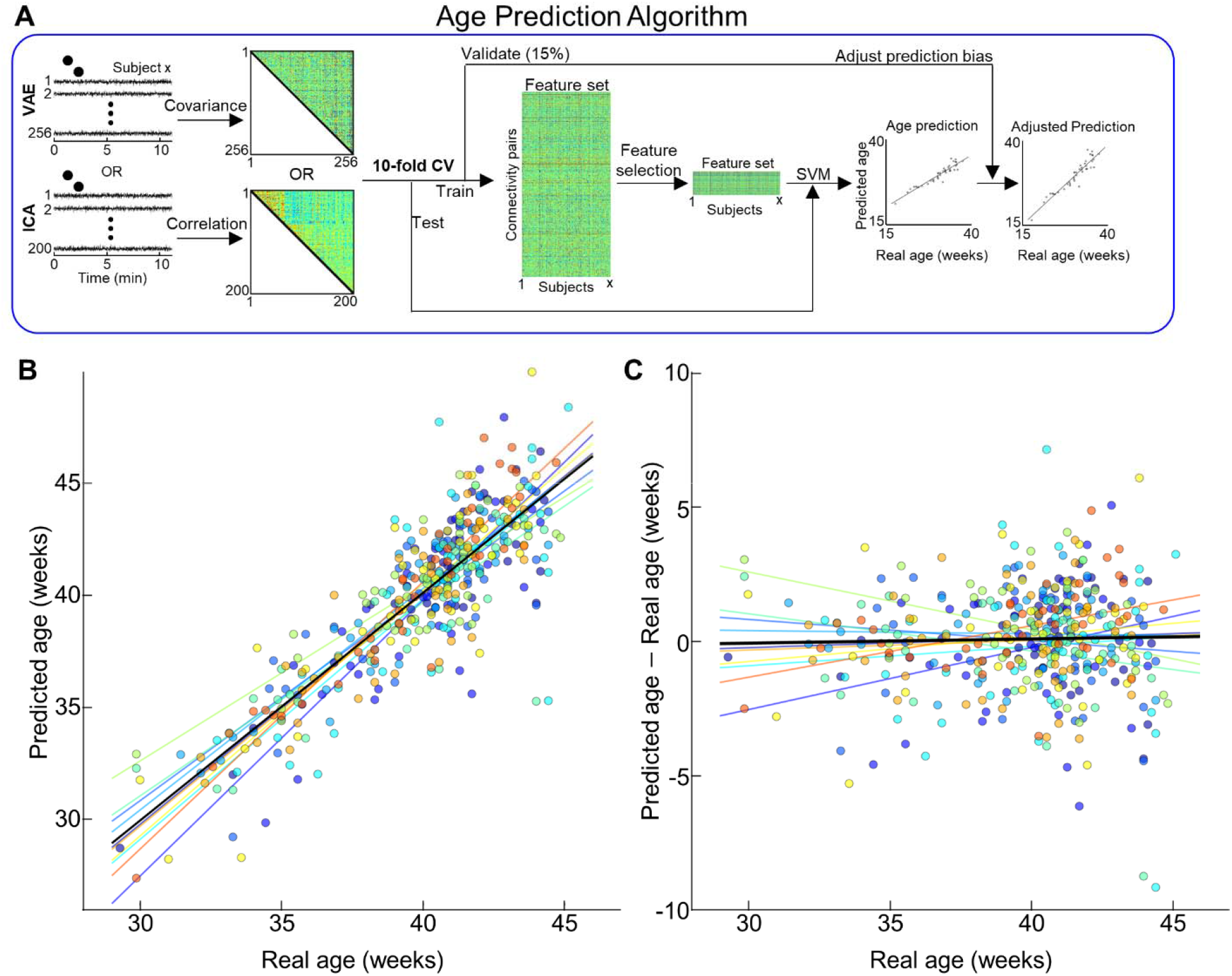
Age prediction based on different latent representations of neonates in the dHCP dataset. **(A)** Illustration of age prediction algorithm. **(B)** Scatterplot between actual age and predicted age. **(C)** Distribution of prediction error across age at scan. **(B, C)** Different colors stand for the prediction age from different folds. Lines with different colors stand for the optimal fit. Black line is the optimal fit for whole samples.

We also tested our age prediction scheme under different latent representations methods in the DBI dataset. Unlike the dHCP dataset, fetuses and neonates in the DBI dataset had fMRI scans with different spatiotemporal resolutions. To minimize possible effects of this difference, we modified the feature selection step in the prediction scheme, deriving the global network strength by summing significant and positive FC edges per subject, yielding a single feature per subject. As the feature space became 1, we substituted linear regression for RSVM. In line with findings in the dHCP dataset, the results clearly showed that latent variables derived by VAE reflected their neurodevelopmental variability across fetuses and neonates more effectively than linear latent representations (Fig. S3; see quantitative result for Table 2). Age prediction in the dHCP dataset was better than the DBI dataset; this is likely due to the shorter scan duration (4-6 mins in DBI dataset vs. 11 mins in dHCP dataset) in the latter.

**Table 2.** Comparison of age prediction performance in DBI dataset using different latent representations. RMSE: root mean squared error, MAE: mean absolute error, R^2^: explained variance. Red highlight indicates the best performance among different latent representation methods.

| Representation Method | RMSE (mean±SD) | MAE | R<sup>2</sup> | Correlation |
| --- | --- | --- | --- | --- |
| VAE (N=256) | 3.59 ± 0.60 | 2.77 ± 0.45 | 0.65 ± 0.10 | 0.78 |
| Cortical Parcel (N=360) | 4.45 ± 1.02 | 3.53 ± 0.83 | 0.56 ± 0.13 | 0.71 |
| IC50 | 5.68 ± 1.72 | 4.49 ± 1.40 | 0.49 ± 0.12 | 0.63 |
| IC100 | 5.14 ± 1.40 | 4.13 ± 1.23 | 0.51 ± 0.14 | 0.68 |
| IC200 | 5.17 ± 1.36 | 4.03 ± 0.90 | 0.51 ± 0.12 | 0.69 |
| IC300 | 4.50 ± 1.04 | 3.56 ± 0.80 | 0.55 ± 0.11 | 0.71 |

### 2.4. VAE-derived representations showed better cross-center generalizability of age prediction than linear representations

Clinical utility of an age prediction model relies on its generalizability across datasets acquired under different recording parameters e.g., spatial and temporal resolution, SNR level etc., For this purpose, we tested the inter-center generalizability of our age prediction model over dHCP and DBI datasets, using different latent spaces. As dHCP and DBI datasets had different recording settings, we re-employed the age prediction model used for the DBI dataset that summed significant and positive FC edges. Here, we used dHCP dataset as training and validation data, and tested the performance of trained models on the DBI dataset, as illustrated in Fig. 5. A. VAE showed the best cross-center prediction performance (Fig. 5. B) compared to linear models (Fig. 5. C-D). Quantitively, we confirmed our observation that VAE showed the least error (RMSE=4.17, MAE=3.37, *r*=0.54), compared to other linear latent spaces (Table 3); cortical parcel and IC maps showed comparable prediction performance. Altogether, these results strongly support our hypothesis that non-linear latent representations featured by the VAE convey neural signatures reflecting brain maturity across a wide range of gestational and postmenstrual ages, in a more effective way than linear latent representations.

**Table 3.** Cross-center generalizability of age prediction performance under different latent representations. RMSE: root mean squared error, MAE: mean absolute error, R^2^: explained variance. Red highlight indicates the best performance among different latent representation methods.

| Representation Method | RMSE | MAE | *r* |
| --- | --- | --- | --- |
| VAE (N=256) | 4.21 | 3.43 | 0.54 |
| Cortical Parcel (N=360) | 8.38 | 7.57 | 0.44 |
| IC50 | 5.05 | 4.04 | 0.17 |
| IC100 | 4.94 | 4.00 | 0.36 |
| IC200 | 5.44 | 4.52 | 0.43 |
| IC300 | 5.28 | 4.42 | 0.44 |

**Figure 5.**
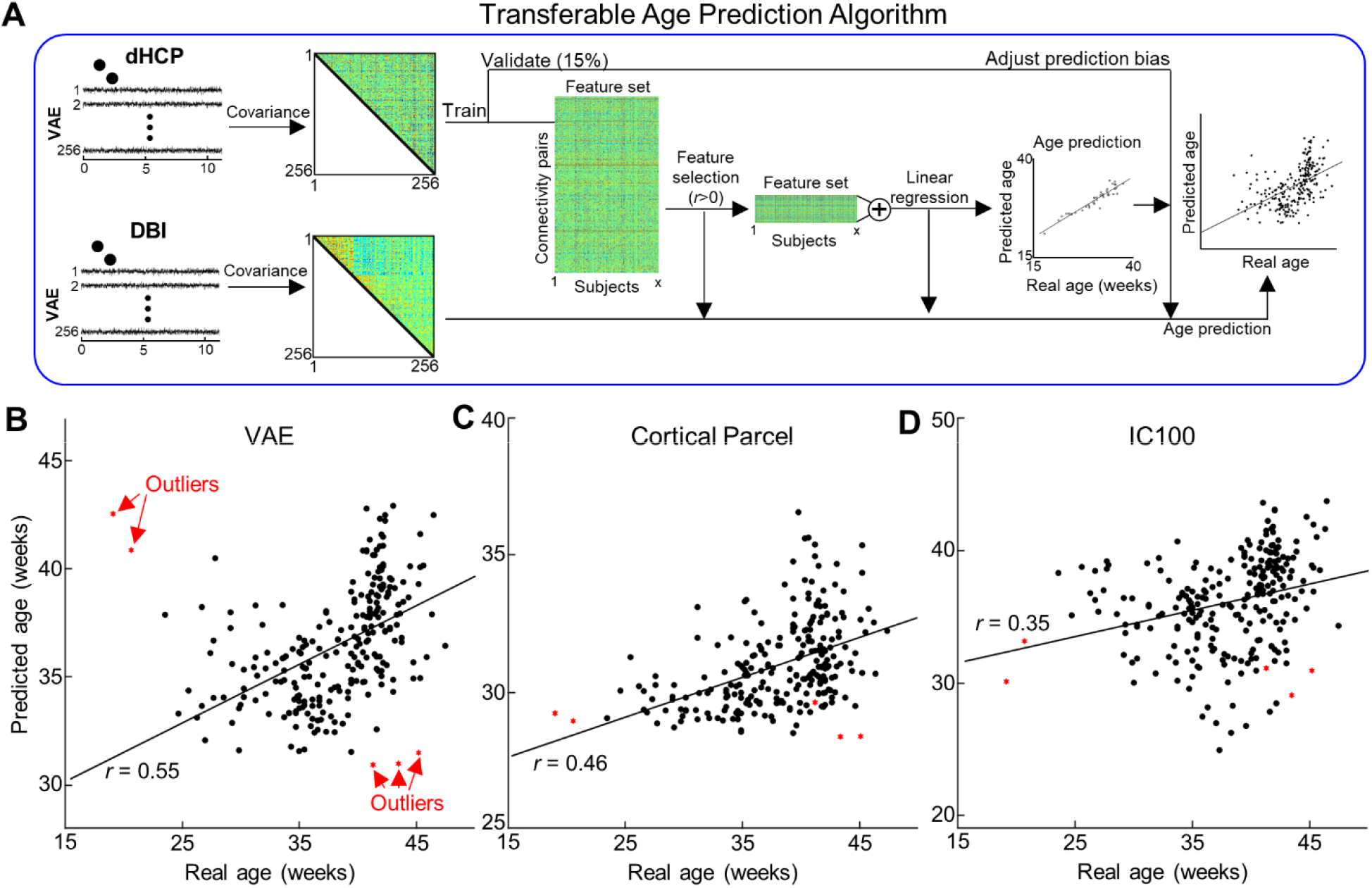
Cross-center age prediction algorithm using different latent representations. **(A)** Illustration of transferable age prediction algorithm. **(B-D)** Scatterplot between actual age and predicted age using different latent representations. Each red dot stands for the outliers having > 3 median absolute deviations away from the median reconstruction performance.

### 2.5. Mapping resting-state brain networks of neonatal babies using VAE

Lastly, we used non-linear latent variables to map functional networks, by utilizing both the VAE encoder and decoder. Briefly, the VAE encoder compressed each timepoint of rsfMRI cortical activity into each latent representation and the VAE decoder visualized latent representations in the cortical surface that we were interested in. As illustrated in Fig. 6. A, per subject, we estimated time-series of latent variables by feeding each brain pattern to the VAE encoder. Estimated latent representations were concatenated across subjects. Then, we re-defined 30 latent bases by applying temporal independent component analysis (ICA) to the concatenated latent variables. Finally, by feeding each independent latent basis into the VAE decoder, we visualized the cortical map of each independent latent basis (see details in Methods). Following this, we investigated functional brain networks (FBNs) for four groups (dHCP vs. DBI; fetus or preterm vs. full-term) separately. For a more optimal comparison between groups, we re-ordered the IC maps based on their pattern similarity by estimating Pearson correlation coefficients between every pair of independent latent bases, each from different groups (center matrix in Fig. 6. B).

**Figure 6.**
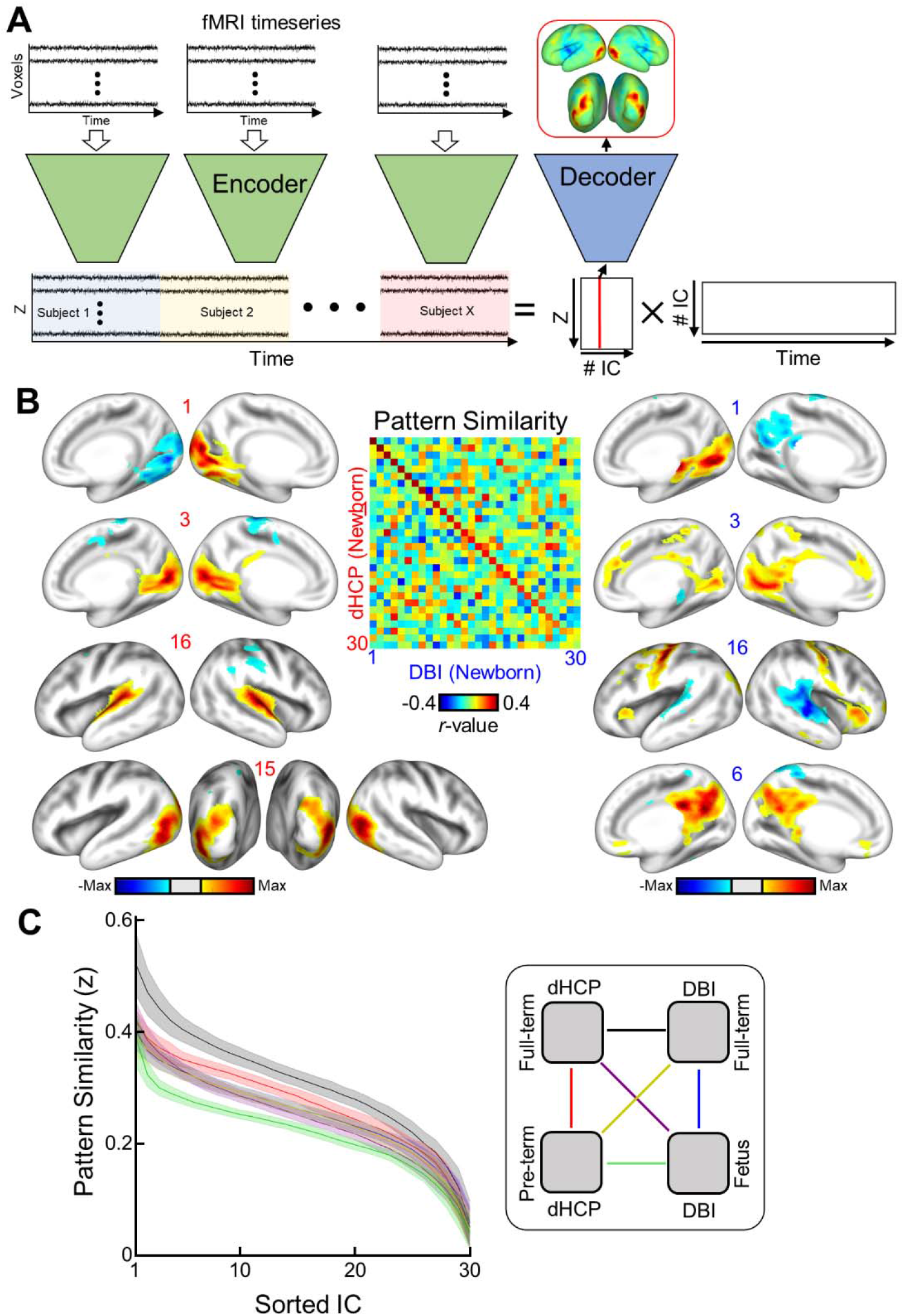
Mapping resting-state functional brain networks of fetuses and neonates using VAE. **(A)** Illustration describing how to map functional brain networks using the VAE. **(B)** Example of neonatal cortical networks estimated from the dHCP dataset (left) or the DBI dataset (right). The order of estimated independent latent variables is sorted by their absolute pattern similarity across different datasets (center). Each map is thresholded at the level of < 15% of maximal absolute value. **(C)** Pattern similarities across different datasets and/or different age groups (coded as lines with different colors; right panel) are plotted. Shades and line stand for the standard deviation and mean similarity across 100 IC results with different initializations.

Each independent latent basis exhibited a unique FBN pattern consisting of both activation and de-activation patterns, as shown in Fig. 6. B. For example, IC1 in both the dHCP and DBI datasets showed activation/de-activation patterns between left and right early visual area whereas IC3 showed only synchronized activation in the primary visual area. This contrast (i.e., activation vs. de-activation) was also observable at the networks level; auditory vs. somatosensory (IC4 in dHCP; IC16 in DBI) and visual vs. auditory (IC16 in dHCP; IC9 in DBI). We also identified the neonatal auditory network having bi-lateral activation in IC16, for both the dHCP and DBI dataset. At the same time, the two datasets showed different brain network patterns. For example, in the dHCP dataset, we found inter-hemispheric opposition of the auditory network (IC20, Fig. S4), and IC15 showed bilateral activation covering the higher visual network (Fig. 6. B left bottom). Conversely, in the DBI dataset, we observed the precursor of the default mode network (IC6, Fig. 6. B right bottom), which coincidently is in-line with findings in young adult rsfMRI studies^19, 34, 35^ global synchrony patterns, which is tightly related to global signal in fMRI data in the dHCP dataset (IC5, Fig. S4), but not in the DBI dataset (Fig. S5). Generally, FBNs identified in the DBI dataset were relatively more complex and less straightforward to interpret, compared to the dHCP dataset, possibly due to the shorter data length in the DBI dataset.

Finally, we tested whether our findings in FBNs were reproducible across different age groups and different datasets. We compared pattern similarity across different datasets and different age groups (Fig. 6. C). The best pattern similarity was achieved when their ages were matched (black line; full-term in dHCP vs. full-term in DBI), rather than between the age-unmatched groups from the same dataset (red line: full-term vs. preterm in dHCP; blue line: full-term vs. fetus in DBI). The least pattern similarity was observed between preterm neonates in the dHCP and fetuses in the DBI dataset. This finding suggests that neurodevelopment of preterm neonates and age-matched in-utero fetuses likely differ given the early extra-uterine exposure in infants born prematurely^36, 37^. Taken together, our results demonstrate that VAE is a valuable tool for investigating early functional brain networks using complex fetal-neonatal rsfMRI data.

## 3. Discussion

Here, for the first time, we used VAE, an unsupervised deep learning model, to extract non-linear representations in human fetal and newborn rsfMRI data. Compared to linear latent space defined by cortical parcels or ICA, non-linear latent space defined by the pretrained VAE was superior in representing rsfMRI patterns acquired using different MRI parameters (Fig. 2). Noteworthy, we found that individual variability in reconstruction performance of brain activity patterns was tightly related to their individual traits that were present, even in the fetal period (Fig. 3). We further demonstrated that non-linear representations extracted by using the VAE provided useful neural signatures of their brain maturity, yielding better age prediction in preterm and term neonates (or fetuses and neonates for the DBI dataset), compared to linear representations (Fig. 4 and S3; Table 1 and 2). By training and testing the prediction model with latent representations across different datasets, dHCP and DBI, respectively, we found that non-linear representations showed the best inter-center generalizability for age predictability (Fig. 5 and Table 3). Finally, we identified 30 functional brain networks in neonates by applying temporal ICA to group-level non-linear representations. Among the 30 brain networks, we identified sensory networks (visual, auditory, somatosensory), precursor of default mode network (Fig. 6; Fig. S4 and S5), and various interaction patterns between brain networks. Collectively, our proposed VAE model and analytical scheme will serve as an important complement to existing linear computational models for disentangling complex fMRI brain patterns in fetuses or neonates for the investigation of longitudinal neurodevelopment in healthy and high-risk fetal-neonatal populations.

In this study, VAE showed better reconstruction performance of fMRI patterns compared to ICA for each subject and across different age groups (Fig. 2. E). Interestingly, VAE outperformed ICA age prediction even when the dimensions of ICA (=300) latent variables exceeded that of the VAE (=256). We believe this stems from the ability of the VAE model to capture non-linear features. In the computer vision field, deep learning studies^38, 39^ have suggested that the non-linear nature of deep convolutional models is key for the model to extract meaningful features against morphological bias/changes of objects, e.g., shift, rotation, or magnification in the image, yielding superior image classification performance. Like morphological distortions in natural images, fMRI images, despite efforts to spatially and temporally denoise the data, may still carry non-neural signals. We posit that a cascade of non-linear operations, efficiently implemented through several layers in our designed VAE architecture, disentangles complex brain activity patterns from residual non-linear noise or artifact, differentiating the VAE model from linear models such as cortical parcel or ICA. Supporting this notion, VAE showed better generalizability of age prediction across different datasets (Fig. 5 and Table. 3).

Recent neuroimaging studies have shown that FC patterns in individuals carry distinct signatures, that are distinguishable, from others^40, 41^. This unique pattern, according to a recent study, remained stable over time for about three years^40^. However, the timing of when these unique individual features emerged is largely unknown. One neonatal study^42^ that explored individual uniqueness of brain FC pattern by identifying individuals based on their FC profiles showed nearly no self-similarity of FC patterns between repeated scans (=11%)^42^, suggesting that the functional uniqueness of the human brain likely emerges after the newborn period. In contrast, another study suggested that individual uniqueness was already apparent during the newborn period reaching an accuracy of 100% (n=40)^43^. However, in this study, data was acquired in a single session, split into two, then used to perform individual identification. This contrasts with other studies that used inter-session similarity to determine uniqueness of functional connectivity profiles. Compared to newborn data^42^, adults (∼92%)^41^, children/youth (6-21 years old, ∼43%)^40^, and infants (∼1 year old, ∼70%)^44^ showed higher inter-scan similarity. In keeping with these data, we observed significant self-similarity – defined as the degree of similarity of reconstructed brain data – between repeated fetal scans (*r*=0.38, Fig. 3. E left) and longitudinal in utero and ex utero scans (*r*=0.39, Fig. 3. E right). As the VAE was trained to represent resting-state brain activity of “typical” adult brain, lower (or higher) reconstruction performance in fetuses or neonates can be interpreted as their lower (higher) similarity of brains representations compared to the adult brain. Future studies investigating the application of reconstruction degree in fetuses and neonates as a potential biomarker of diseases, or a predictor of emotional/behavior outcome will be of interest.

In adults^29^, we hypothesized that our trained VAE model was able to extract generative factors of rsfMRI data as a form of latent variable. In the VAE, we speculated that learned generative factors, i.e., latent variables, largely fall into two groups: one representing the common pattern of brain dynamics and the other shaping the unique brain patterns of individuals. Supporting this notion, in the current study, we observed that a subset of latent variables of fetal-neonatal rsfMRI mapped to a set of large-scale functional brain networks e.g., visual, auditory, and sensorimotor, and default mode network (Fig. 6; Fig. S4 and S5), whereas another subset reflected the temporal properties of fetal-neonatal functional brain development (Fig. 4-5; Fig. S3; Table 1-3). Our findings suggest the delineation of generative factors of fetal-neonatal rsfMRI is plausible using the VAE model pretrained on adult rsfMRI without any further introduction of new fetal-neonatal fMRI data to the VAE training. Collectively, we believe that VAE may be used to create longitudinal maps of the functional brain connectome throughout the lifespan (i.e., a functional atlas from the fetal period to adulthood).

Central to this study was the improvement in spatial representation of fetal-neonatal fMRI cortical patterns via the VAE pretrained from resting-state brain patterns of adults. Once the fetal-neonatal rsfMRI data were projected onto the latent space defined by the VAE, we were able to model the temporal dynamics of latent representations at the group level using linear computational methods e.g., temporal ICA. To this end, we identified 30 independent latent bases and visualized each of them through the VAE decoder (Fig. 6). Compared to functional brain networks defined by spatial ICA on the dHCP dataset^17^, brain networks reported in our study exhibited more dynamic patterns such as lateralized activation, bilateral activation, activation and deactivation across hemisphere, and interaction between brain networks (Fig. 6; Fig. S4 and S5). We believe the discrepancy in functional brain network patterns mainly originates from the methodological difference between temporal and spatial ICA. As shown in previous studies with adults^45^, different from spatial ICA, temporal ICA outputs temporally-independent functional modes having activation patterns along with deactivation patterns. Similarly, we also observed the opposition between sensory-related networks (#4: auditory vs. somatosensory; #16: visual vs. auditory, Fig. S4). However, unlike the adult brain, we were not able to observe anticorrelated activity between cognitive networks and sensory networks, possibly suggesting such suppression of sensory functions via higher cognitive networks develops at a later neurodevelopmental phase. We expect our proposed analysis scheme combining spatial compression via VAE and temporal modeling via temporal ICA may be a critical addition to existing tools for mapping fetal-neonatal brain networks.

While the VAE model ably represented fetal-neonatal FC data, one limitation is that the model primarily focused on learning spatial representations of rsfMRI patterns and ignored the temporal dynamics of rsfMRI data. Functional MRI is not stationary^46^ and our future work aims to address both spatial and temporal features of rsfMRI; currently, limitations on technical resources constrained our ability to simultaneously model both. One recent study done by Qiang *et al*.,^47^ tried to address this issue by introducing recurrent neural network to the VAE model. However, their model had to sacrifice the efficacy of their spatial representations, by diminishing the geometrical structure of fMRI data (from 3D image +1D temporal to 1D image +1D temporal). Another recent study^48^ was able to keep the 3D structure of fMRI data, but down-sampled the spatial resolution of input image grossly and reduced the computational layers, for the sake of computational resources. Both models have achieved their desired temporal interpretability but at the cost of degraded spatial representations. Future studies with novel designs are needed to model the complex and multifaceted spatiotemporal dynamics of fMRI data.

In this study, we applied a novel deep generative model to a large (>500) fetal-neonatal rsfMRI dataset to extract non-linear brain representations of healthy fetuses, preterm neonates and full-term infants. Compared to linear methods, the VAE model generated improved representations of the brain’s functional connectivity patterns and a more precise age prediction. Characterization of the developing brain’s functional connectome using the VAE model provided important insights into the complex fetal-neonatal neurodevelopmental growth trajectories that were previously inaccessible using conventional linear models. These observations have provided important insight into the complex non-linear brain development that occurs during the prenatal-neonatal continuum that may allow us to identify early biomarkers for neurobehavioral vulnerability and link specific prenatal brain developmental processes associated with aberrant neural connectivity which may underly prevalent child and adult neurologic disorders.

## 4. Materials and Methods

We analyzed two large fMRI datasets (N=727 scans), one dataset consisting of fetuses and neonates collected by our institute (n=270), named as “Developing Brain Institute” or DBI dataset, and one public dataset consisting of pre- and full-term neonates (n=457), published by developing human connectome project, named as dHCP dataset^49^. Two fetal-neonatal datasets (DBI and dHCP), acquired by two independent centers, were utilized to test the utility and generalizability of the VAE model on representing fetal-neonatal rsfMRI data.

### Developing Brain Institute (DBI) dataset: fetus and full-term neonate

We analyzed 270 resting state scans from 95 healthy fetuses and 160 full-term born healthy infants. Scans were collected as a part of an ongoing longitudinal study examining prenatal-neonatal brain development at Children’s National Hospital in Washington DC. Fetuses (49 female) from healthy pregnancies were scanned between 19.14 – 39.71 gestational weeks (mean and SD; 33.65±4.01). Postmenstrual age of full-term born infants at scan ranges between 37.71 – 47.43 weeks (41.67±1.84). Maternal exclusion criteria were psychiatric disorders, metabolic disorders, genetic disorders, complicated pregnancies, multiple pregnancies, alcohol, and tobacco use, maternal medications, and contraindications to MRI. All experiments were conducted under the regulations and guidelines approved by the Institutional Review Board (IRB) of Children’s National; written informed consent was obtained from each pregnant woman who participated in the study.

For fetal scans, structural and functional resting-state MR images were acquired using a 1.5 Tesla GE MRI scanner with 8-channel receiver coil. The structural MR images for the fetal brain were acquired using single-shot fast spin-echo T2-weighted images by following settings: TR=1100ms, TE=160ms, flip angle=90 degrees, and voxel size= 0.8×0.8×2mm. Functional data were acquired using echo planar images (EPI) with TR=3000ms, TE=60ms, flip angle=90 degrees, field of view= 33cm, and voxel size = 2.58×2.58×3mm, and total scan volume=144 (=7.2min). The structural and functional MR images of full-term infant brain MRI studies were acquired using 3T GE scanner. T2-weighted fast spin echo MRI was obtained using the following parameters: TR=2500ms; TE=64.49ms, voxel size= 0.625×1×0.625mm. The parameters of fMRI MRI scans were set to TR=2000ms, TE=35ms, voxel size=3.125×3.125×3mm, flip angle= 60 degrees, field of view=100mm, and total scan volume=200∼300 (=6.7∼10min).

Functional MR images were preprocessed as followings: slice time correction, bias-field correction, motion-correction, spatial smoothing at full-width-half-maximum=4.5mm, and intensity scaling. We excluded volumes having excessive head motion (frame-wise motion >1mm or rotational motion >1.5 degrees). Details of the preprocessing steps for the fetal MRI data can be found at^4^ and details regarding neonatal scans can be found at ^50^. Neonatal MRI scans with < 4 mins were excluded in the analysis. The analyzed data length of fetus and neonate group was 4-7 (mean±S.D.=5.4±0.9) and 4-8.9 (5.5±0.8) mins, respectively. Finally, preprocessed rsfMRI scans at the volumetric brain space were projected to the standard cortical space using HCP workbench command *-volume-to-surface-mapping*.

### Developing Human Connectome Project (dHCP) dataset: preterm and full-term neonates

Developing Human Connectome Project (dHCP) is an open science project approved by the UK National Research Ethics Authority^49^. FMRI data analyzed in the project can be downloaded at https://data.developingconnectome.org/. Among 464 subjects (261F/203M), we analyzed a total of 457 resting state fMRI (rsfMRI) scans from 107 preterm and 302 full-term neonates. Their postmenstrual ages at scan range from 24.29 weeks to 44.87 weeks, and the mean and standard deviation of ages were 38.54 and 3.47 weeks, respectively. It was reported that no babies in the preterm and full-term group had major brain injury. Demographical details regarding the dataset can be found elsewhere at ^38^. MR scans were acquired using a 3 T Philips Achieva system (Philips Medical Systems). High-resolution T1- and T2-weighted anatomical imaging was acquired with followed parameter settings: T1-weighted image; spatial resolution= 0.8mm isotropic, FOV= 145×122×100mm, and TR=4795 ms, T2-weighted image; spatial resolution= 0.8mm isotropic, FOV= 145×145×108mm, TR=12000ms, and TE=156ms. Followed by anatomical scans, fMRI scans were acquired with acquisition specifications: multislice gradient-echo echo planar imaging (EPI) sequence, multiband factor=9. TR=392ms, TE=38ms, flip angle=34, spatial resolution= 2.15 mm isotropic, and total volume=2300 (∼15 mins). The downloaded dHCP fMRI data was already preprocessed^51^ and registered into the volumetric brain template^52^. To match the input format to the required input format of the VAE model, we wrapped volumetric fMRI data to the cortical space using HCP workbench command *-volume-to-surface-mapping*. After it, we applied additional preprocessing step consisting of voxel-wise detrending (regressing out a 3rd-order polynomial function), bandpass filtering (from 0.01 to 0.1 Hz), and voxel-wise normalization (zero mean and unitary variance). To avoid possible distortion from the filtering step, we rejected first and last 150 volumes from the analysis. Lastly, to prevent the possible confounding effects from motion artifact, we selected 1,400 volumes (∼10 mins) having the least frame-wise displacement degree, per subject.

### Pretrained Variational Autoencoder using Adult rsfMRI

In our previous work ^29^, we designed a variation of fJ-VAE^53, 54^, to learn representations of rsfMRI patterns of adults. Briefly, our proposed VAE model consisted of an encoder and a decoder. After reformatting the fMRI pattern at the cortical space to the carefully designed 2-D regular grid space (Fig. 1. A), an encoder compressed an fMRI map (# of vertices=38,864) to a probabilistic distribution of 256 latent variables, whereas a decoder reconstructed the fMRI patterns given the sampled latent variables (Fig. 1. B and C). The encoder and decoder consisted of 5 non-linear convolutional layers and 5 de-convolutional layers, respectively. The overarching goal of the VAE model was to reconstruct input at best under the constraint forcing the distribution of every latent variable to be close to be i.i.d. Specifically, given the training dataset X, we optimized the encoding parameter *ϕ* and the decoder parameter *θ*, to minimize the loss function defined as:

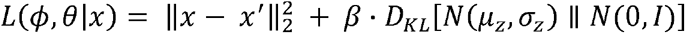

where x and x′ are the input image and the reconstructed version of input image, *β* is a hyperparameter balancing the two terms in the loss function, *D*_*KL*_ is the Kullback-Leibler (K-L) divergence between the posterior distribution *N*(*µ*_*z*_, *σ*_*z*_) and prior distribution *N*(0, *I*), and *µ*_*z*_ and *σ*_*z*_ are mean and standard deviation of estimated latent variables z from x. Upon validation dataset consisting of 50 young adults rsfMRI data from Human Connectome Project ^32^, we optimized the hyperparameters of VAE model: total number layers=12, learning rate=10^−4^, batch size=128, and *β*=9. After being trained with another large HCP rs-fMRI data consisting of 100 young adults, the trained VAE model has learned to represent patterns of cortical activity and network patterns using latent variables. Pleases refer to our original paper ^29^ for the detailed description of geometric reformatting and the model architecture of the VAE. As our goal of this study was to utilize brain representations learnt from adults for better understanding of fetal-neonatal neuroimaging data, we froze the model parameters of the VAE model, *ϕ* and *θ*, and froze the VAE model parameters. Therefore, the VAE model used in this study was identical to the VAE model reported in our original paper^29^. In this study, the dHCP and DBI datasets were only used for testing the pretrained VAE model. Latent variables estimated from cortical patterns of the dHCP and DBI datasets were used as features of age prediction experiment or networking mapping (Fig. 1. D).

### Reconstructing fetal-neonatal rsfMRI patterns using pretrained VAE

We evaluated how well the VAE model that was pretrained to represent adult rsfMRI delineates the fetal-neonatal brain activity that was unseen during the training. We compressed the spontaneous cortical patterns of the dHCP and DBI datasets using the pretrained VAE encoder and restored them to the cortical space by feeding sampled latent variable to the VAE decoder. The reconstruction degree of each time point was defined as the Pearson correlation coefficient between the original- and reconstructed cortical patterns. Averaging reconstruction performance over timeseries or subjects was done after converting *r*-value into Fisher’s *z*-score.

### Predicting gestational age and postmenstrual age of fetus, pre-term neonates, and full-term neonates

To evaluate the age prediction power of latent variables defined by the VAE, we built an age prediction model and validated its predictability using 10-fold cross validation scheme. For dHCP dataset, we utilized FC profile at the latent space as individual-wise features (Fig. 4. A). Specifically, cortical pattern of each time point was encoded as timeseries of latent variables using the VAE encoder. Next, we measured FC profile (# of features=256×256/2) of each subject by measuring covariance between every pair of timeseries of latent variables. After keeping 10% of dataset as a testing group, we further divided the remainder 90% of dataset into training (=76.5%) and validation set (=13.5%). Within the training dataset, we applied the feature selection procedure by correlating the age and each feature. We thresholded features having FDR-corrected *q* < 0.05. Based on the selected feature set, we built a linear regression support vector model (RSVM) using MATLAB function *fitrsvm*.*m*. In comparison to other age prediction studies ^33^, our regression model also tended to output biased error towards extremity of age distribution, called as a prediction bias. To eliminate such prediction bias, we further added an adjusting step of predicted age using the validation dataset. Specifically, we predicted the age of validation dataset using the trained RSVM model and built a linear regression model between predicted- and actual age of validation set. After calculating the optimal regression fit, we compensated the prediction age by the trained RSVM model. Finally, the prediction performance of designed model was tested using the unseen testing dataset, repeating the whole procedure for 10 folds independently.

The prediction model for the DBI dataset was nearly identical to the above prediction model in dHCP dataset, except that, in the feature selection step, we took the feature positively correlated to the age and summed the significant features. The reason behind this modification was to make the model more robust against different recording specifications (e.g., spatial resolution) between fetal scans and neonatal scans in the DBI dataset. As the dimension of feature set became 1, we employed a linear regression model using MATLAB function *fitlm*.*m*. Note that, with the exception of the feature selection step and regression model type, all other steps remained identical as the one used in the dHCP dataset.

Finally, we built an age prediction model generalizable across different datasets, dHCP and DBI (Fig. 5. A). We took the whole dHCP dataset as training/validation set for the model and tested the prediction performance on the DBI dataset. We separated the dHCP dataset into testing set (=85%) and validation set (=15%). The age prediction algorithm was identical to one employed in the DBI dataset, as two datasets had different spatial and temporal resolution (e.g., 10 mins for dHCP vs. 4∼6 mins for DBI).

The prediction performance was evaluated using four different metrics, root-mean-squared-error (RMSE), mean-absolute-error (MAE), squared correlation coefficient per fold (*r*^2^), and correlation coefficient between actual age and predicted age at the whole dataset level (correlation).

### Mapping of fetal-neonatal brain networks using pretrained VAE

We examined the utility of latent variables defined by the VAE as a mapping tool of fetal-neonatal functional brain networks (Fig. 6. A). First, we estimated the timeseries of latent variables by feeding each fMRI time point to the VAE encoder, and concatenated the timeseries of latent variables across subjects, for different age groups; pre-term baby (<27weeks) vs. full-term neonates (>=27weeks) for dHCP dataset and fetuses vs. full-term neonates from the DBI dataset. We re-defined the latent space by applying temporal independent component analysis (ICA) to concatenated latent variables. The temporal ICA was applied using the Infomax ICA algorithm, which was implemented by MATLAB toolbox EEGLAB^55^. Inspired by the recent ICA study investigating neonatal functional brain networks^17^, we also set the number of ICs to 30 heuristically. Once 30 independent latent bases were identified, cortical mapping of each was done as followings: 1) we multiplied random scaling factors (n=1,000) to the latent basis, 2) reconstructed 1,000 cortical patterns of scaled latent basis using the VAE decoder, and 3) calculated covariance between random scaling factors and reconstructed fMRI activity, per cortical location. The estimated covariance scale of each cortical location was considered as the activation level of the location and the sign (+/-) of covariance was defined as activation (+) or deactivation (-) state. We fixed the sign of each cortical map as the region having the strongest covariance scale being activation (+) state. Note that as the VAE decoder is highly non-linear, our proposed mapping method from latent space to cortical space did not guarantee to reflect the true cortical meaning of independent latent bases, but the results in our study empirically suggested that our proposed method was a good approximation mapping strategy. Lastly, the cortical map of each basis function was thresholded at the 15% of the maximal absolute value of each map, for the better interpretation of results.

We further examined group-wise reproducibility of estimated independent latent bases across age groups and/or different datasets. Similarity between latent bases was measured by estimating the Pearson correlation between every latent basis from one group and every basis from another group. As the sign of independent latent bases was somewhat arbitrarily determined in ICA algorithm, we one-to-one paired latent bases across age groups based on their absolute correlation coefficient values. We took the trend of paired similarity over increasing IC numbers as the inter-group reproducibility of brain network. For the sake of reproducibility, the whole analysis was independently repeated for 100 times, each with different initialization seeds for ICA. The final reproducibility trend was reported by averaging over 100 trials.

### Comparison with linear representation methods

To access the utility of the VAE model, we employed conventional linear representation methods, cortical parcellations and group ICA maps. For the sake of fair comparison between representation methods, we utilized multi-modal cortical parcellation^56^ and group ICA maps^57^, both were derived by the HCP dataset. In the employed cortical parcellation, there were 360 areas (180 per hemisphere), and we compressed fMRI data at the cortical space into the linear latent space (# of dimension=360 cortical parcels). We also utilized group ICA results provided by HCP, which is freely available at https://db.humanconnectome.org/. Briefly, group ICA maps were estimated based on the rsfMRI of 820 subjects in the HCP and the complete algorithm of group ICA can be found at^32^. This group ICA provides different number of latent variables (or # of IC), 15, 25, 50, 100, 200, and 300. Among them, we utilized IC maps consisting of 50, 100, 200, and 300 ICs (named as IC50, IC100, IC200, and IC300). Similar to VAE model, we compressed cortical activities into IC maps and restored them using the pseudoinverse of IC maps. Same as VAE model, the reconstruction performance was measured by correlating between original- and reconstructed cortical patterns. Linear latent variables defined by different IC maps were utilized for age prediction task as well.

## Acknowledgments

We thank the participants of this study. This study was funded by grant R01 HL116585– 01 from the National Heart, Lung, and Blood Institute, National Institutes of Health and grant MOP-81116 from the Canadian Institute of Health Research.

## Data and materials availability

The data and code that support the findings of this study are available from the corresponding author, CL, upon reasonable request.

